# General cognitive performance declines with female age and is negatively related to fledging success in a wild bird

**DOI:** 10.1101/2022.08.30.505947

**Authors:** Camilla Soravia, Benjamin J. Ashton, Alex Thornton, Amanda R. Ridley

## Abstract

Identifying the causes and fitness consequences of intraspecific variation in cognitive performance is fundamental to understand how cognition evolves. Selection may act on different cognitive traits separately or jointly as part of the general cognitive performance of the individual. To date, few studies have examined simultaneously whether individual cognitive performance covaries across different cognitive tasks, the relative importance of individual and social attributes in determining cognitive variation, and its fitness consequences in the wild. Here, we tested 38 wild southern pied babblers (*Turdoides bicolor*) on a cognitive test battery targeting associative learning, reversal learning and inhibitory control. We found that a single factor explained 59.5% of the variation in individual cognitive performance across tasks, suggestive of a general cognitive factor. General cognitive performance varied by age and sex; declining with age in females but not males. Older females also tended to produce a higher average number of fledglings per year compared to younger females. Analysing over 10 years of breeding data, we found that individuals with lower general cognitive performance produced more fledglings per year. Collectively, our findings support the existence of a trade-off between cognitive performance and reproductive success in a wild bird.

## 1. INTRODUCTION

The mental mechanisms through which animals acquire, process, store and act on information from the environment represent animal cognition [1]. Animals use cognitive mechanisms to adjust their behavioural responses to the environmental and social context, remember the location of resources, and learn which environmental cues indicate presence of food, mates or predators [2]. Different animal species rely on cognitive mechanisms to different extents to solve ecological problems [e.g. 3, 4], and even within a species, cognitive performance can vary significantly among individuals [e.g. 5, 6]. Such great inter-and intraspecific variation has led to the question: what selective pressures shape the evolution of cognition?

Intraspecific studies of animal cognition have shown that in some cases cognitive performance is heritable [7]. Additionally, cognitive performance has been linked to mate choice [8], reproductive investment [6], reproductive success [5, 9], and survival [10]. The existence of differential fitness arising from heritable variation in cognitive performance means cognitive traits can evolve [11]. However, better cognitive performance is not always associated with increased fitness, for example, faster learning is associated with reduced longevity in fruit flies (*Drosophila melanogaster*) [12]. This may occur because the energetic costs of enhanced cognitive function lead to a trade-off between resource allocation to cognitive performance and somatic maintenance or reproduction [11, 13]. Therefore, we expect selection to favour cognitive performance only when the benefits outweigh the costs [13].

To understand how selection acts on cognition, the link between cognitive variation and fitness consequences needs to be identified, as well as the proximate causes of individual variation in cognitive performance. Several factors have been associated with differences in individual cognitive performance, including the physical and social environment [14, 15], and individual attributes such as age [e.g. 16], rank [e.g. 17], and sex [e.g. 18]. Sex differences in cognitive performance often arise as a consequence of mating strategies or sex-specific ecological constraints [19, 20]. For example, in the brood-parasitic brown headed cowbird (*Molothrus ater*), females outperform males on a large-scale spatial memory task, likely because the breeding strategy of this species relies on females finding potential host nests to lay their eggs in [21].

Age differences in cognitive performance have been mostly found when comparing juveniles and adults [22, 23]. However, cognitive performance can also change during adult life [24, 25]. The gradual reduction of cognitive function with age is known as cognitive senescence [26]. Evidence for cognitive senescence in non-human animals is largely limited to captive studies [24, 27]. For example, homing pigeons (*Columba livia*) older than 10 years returned more often to feeders that they had just depleted despite them being empty, showing impaired short-term memory [16]. To date, little is known about cognitive senescence in the wild, due to logistical limitations including difficulties of estimating individuals’ age [28] and testing cognition in the wild [29].

During social interactions, individuals can differ in rank, where dominant individuals often monopolize resources [30]. There is growing evidence that cognitive performance is related to rank, but the direction of this relationship varies across studies [31, 32]. It has been suggested that cognitive performance may not be related to social status *per se*, but to factors correlated to rank, such as vigilance, neophobia or motivation to find alternative food sources [17, 32]. For example, in Arabian babblers (*Argya squamiceps*) subordinates were the first to learn to remove black lids in a novel foraging task, likely because they were more explorative, but dominants, which tend to be older in this species, were better able to generalise the solution to white lids because of experience [33, 34]. Finally, individual cognitive performance may also be linked to social group size because living in larger groups may require better cognitive performance in order to monitor the state and actions of group members, remember their identity, and the outcome of past interactions [5, 14]. Despite the growing number of studies investigating intraspecific differences in cognitive performance, these individual and social attributes (rank, sex, age and group size) have rarely been examined simultaneously while controlling for proxies of motivation, and evidence of their relative importance in driving cognitive variation is scarce.

If we identify what selection pressures drive variation in cognition, the question of whether these act on each cognitive trait separately, or jointly as part of general cognitive processes, remains. In humans, it has been repeatedly demonstrated that individual performance correlates positively across different cognitive tasks, and approximately 40% of the total variation in performance can be explained by a single general cognitive factor *g* [35]. This factor, also referred to as general intelligence or intelligence quotient (IQ), predicts important life outcomes, such as occupational attainment, health and longevity [36]. Recently, several studies in non-human animals have also described something akin to a general cognitive factor *g* explaining between 30%-60% of variation in cognitive performance across a battery of cognitive tasks [reviewed in 37]. The evidence for *g* provided by animal studies however has encountered criticism. First, generating reliable measures of *g* in non-human animals requires the use of robust psychometric test batteries targeting well-studied cognitive traits [38]. It is also worth noting that variation in the combination of cognitive tasks used in a test battery can lead to different estimates of *g* [39]. Second, results indicative of *g* may also arise in the absence of a truly general cognitive factor if performance on different tasks is underpinned by the same cognitive mechanism – for instance, variation in associative learning performance could potentially impact performance across a range of tasks [40]. Therefore, the single factor extracted from animal cognitive test batteries does not necessarily equate to general intelligence or *g* as described in humans. Nonetheless, if performance measured across a battery of cognitive tasks can be explained by a single factor, hereafter referred to as “general cognitive performance (GCP)” [5, 41], and this factor predicts fitness in the wild [5], then it may represent a measurable cognitive trait which may be under selection in animal populations [35].

Here, we tested wild adult southern pied babblers (hereafter “babblers”, *Turdoides bicolor*) on a psychometric test battery containing three tasks designed to quantify (1) associative learning, (2) reversal learning, (3) inhibitory control. These are well-studied cognitive traits that span different domains [38, 42]. Additionally, they are likely to be ecologically relevant as they allow individuals to: learn predictive contingencies between environmental cues (associative learning); learn a new association when the previous one stops being rewarding (reversal learning); and control prepotent motor responses when counterproductive (inhibitory control) [42, 43]. To achieve a comprehensive understanding of the relationship between different cognitive traits, the factors underpinning interindividual variation in cognition and the link between cognitive performance and fitness in a wild animal population, we: (a) tested whether individual cognitive performance was positively correlated across tasks and could be explained by a single factor (GCP); (b) measured proxies of motivation and attributes of the individual and social group (age, sex, rank, group size) to identify determinants of individual cognitive performance; and (c) related individual cognitive performance to multiple measures of reproductive success.

## 2. METHODS

### 2.1 Study site and species

Data were collected at the Kuruman River Reserve (26°58′ S, 21°49′ E; South Africa, 33 km^2^) between September-March in 2018, 2019 and 2021. The reserve is situated within the semi-arid Kalahari region, which is characterized by vegetated sand dunes [44]. Pied babblers are medium-sized (60-90 g), sexually monomorphic passerines endemic to this region. They are cooperative breeders and live in groups, which include a dominant breeding pair and subordinate helpers [45]. The dominant pair produces approximately 95% of the offspring [46, 47]. Pair bond tenure varies greatly (from < 1 month to > 5 years) [48]. All adult group members (> 1 year post-hatching) engage in care of young and territory defence [45]. Each group defends a territory of 50-80 hectares year-round [49]. On average only 4% of subordinates live in non-natal groups each year [46].

The study population has been monitored since 2003 and is habituated to human presence [45], which allows researchers to observe the birds’ natural behaviour from a close distance (< 5 m) and to present them with cognitive tasks. Ringing and blood sampling for molecular sexing are performed on nestlings 11 days post-hatching [50]. Therefore, each bird in the study population is identifiable by a unique ring combination, and sex and age are known for all adult birds. Adult immigrants are trapped with a walk-in trap for ringing and blood sampling. We considered immigrants to be at least one year old at the time they immigrated into our study population, and if they immigrated as dominants and bred on the first year in which they immigrated we considered them to be at least two years old, as dispersal and first breeding are rarely recorded before these ages respectively [48, 51]. On average, subordinate individuals are younger than dominants [51]. Rank (dominant vs subordinate) is easily inferred from aggressive displays by the dominant individuals towards subordinates [45], and distinctive affiliative behaviours between dominants [48]; in addition, only the dominant female incubates the nest overnight [45]. During the study years (2018-2021), the population comprised 14 groups ranging in size from two to seven adults. We tested different individuals each year: 13 individuals from six groups in 2018, 18 from 10 groups in 2019 and seven from four groups in 2021. Among the birds tested in 2021, four were unringed when tested because we found them as yearlings after a year’s gap in data collection (fieldwork in 2020 was suspended due to the COVID-19 outbreak). We were able to identify these birds based on distinctive individual features (e.g. plumage or scarring), but their sex was unknown. Sex was unknown also for two other ringed individuals tested in 2021 due to delays in the analysis of blood samples caused by COVID-19.

### 2.2 Cognitive test battery

The cognitive test battery consisted of three tasks designed to quantify (1) associative learning, (2) reversal learning, (3) inhibitory control. These cognitive tasks tapped into the natural terrestrial foraging behaviour of babblers [45], as they required them to peck downwards at a lid or move around a barrier on the ground to retrieve a food reward: a mealworm (*Tenebrio molitor* larva). The original cognitive test battery included a spatial memory task, but this was later excluded because individuals’ behaviour when interacting with the task did not deviate from a random sampling strategy (see Supplementary Material section 3).

Cognitive testing was conducted between 5 am and 7 pm, when babblers were active. Cognitive tasks were always presented in the shade when the birds were not showing any heat dissipation behaviours (i.e. panting and wingspreading) to avoid potential confounding effects of heat stress on cognitive performance [52]. All trials in a cognitive test were performed when the focal individual was temporarily out of sight of other group members. This was achievable because of the short trial duration (< 1 min) and because babblers often forage over 10 m apart from each other [53]. The three cognitive tests were carried out at least 24h apart and the order was randomised within individual, except for the reversal learning, which was always carried out the day after the associative learning test. Prior to quantifying learning performance, individuals were trained to peck the lids in a cognitive task to find a food reward using unpainted lids (see Supplementary Material section 1). In all tasks, if the focal bird did not interact with the task for 30 min, the test was paused and continued the following day, and if the passing criterion was not reached by 120 trials, the test was stopped.

#### 2.2.1 Associative and reversal learning

The task used to quantify associative and reversal learning consisted of a small wooden block (180 × 70 × 30 mm) with two equidistant circular wells (30 mm diameter, 20 mm depth) covered by painted wooden lids. The lids were held in place by elastic bands; in this way, they fitted snugly into the wells, preventing the bird from using visual cues to identify the rewarded well, but they could swivel when pecked, making the food reward accessible to the bird (Figure 1A). The two lids were painted a dark and light shade of the same colour rather than two different colours to avoid effects of past experience or colour salience on learning performance [e.g. 29, 41; hereafter “colours” instead of “colour shades” for brevity]. Each day before the start of cognitive testing, two mealworms were temporarily placed in both wells of the cognitive task in order to prevent the bird from relying on olfactory cues to choose the rewarded well during testing. The associative and reversal learning tests followed the protocols used by Shaw et al. [41] and Ashton et al. [5]. One of the two colours was randomly assigned to be the rewarded colour for each test bird. In each trial, the first peck of the individual when approaching the task was counted as correct (1 = rewarded lid) or incorrect (0 = unrewarded lid). During the first trial, the individual was allowed to search both wells to see that only one hid the reward. In the following trials, if the individual chose correctly, it ate the mealworm and the task was removed to replace it out of its sight. If the individual chose incorrectly the task was removed before the individual could peck the other lid and gain the reward. The position of the rewarded well was pseudorandomised between trials to ensure the individual associated the colour of the lid with the reward, and not the position of the lid. Associative learning performance was quantified as the number of trials required to reach the passing criterion, which was six correct choices in a row (a significant deviation from a random binomial probability: binomial test p = 0.016; i.e. the individual has a probability of 1.6% of achieving six correct choices in a row by random chance). If the bird passed the associative learning task, the reversal learning task was carried out 24h after. Reversal learning performance was quantified using exactly the same protocol and passing criterion used for associative learning, but rewarding the opposite colour.

**Figure 1.**
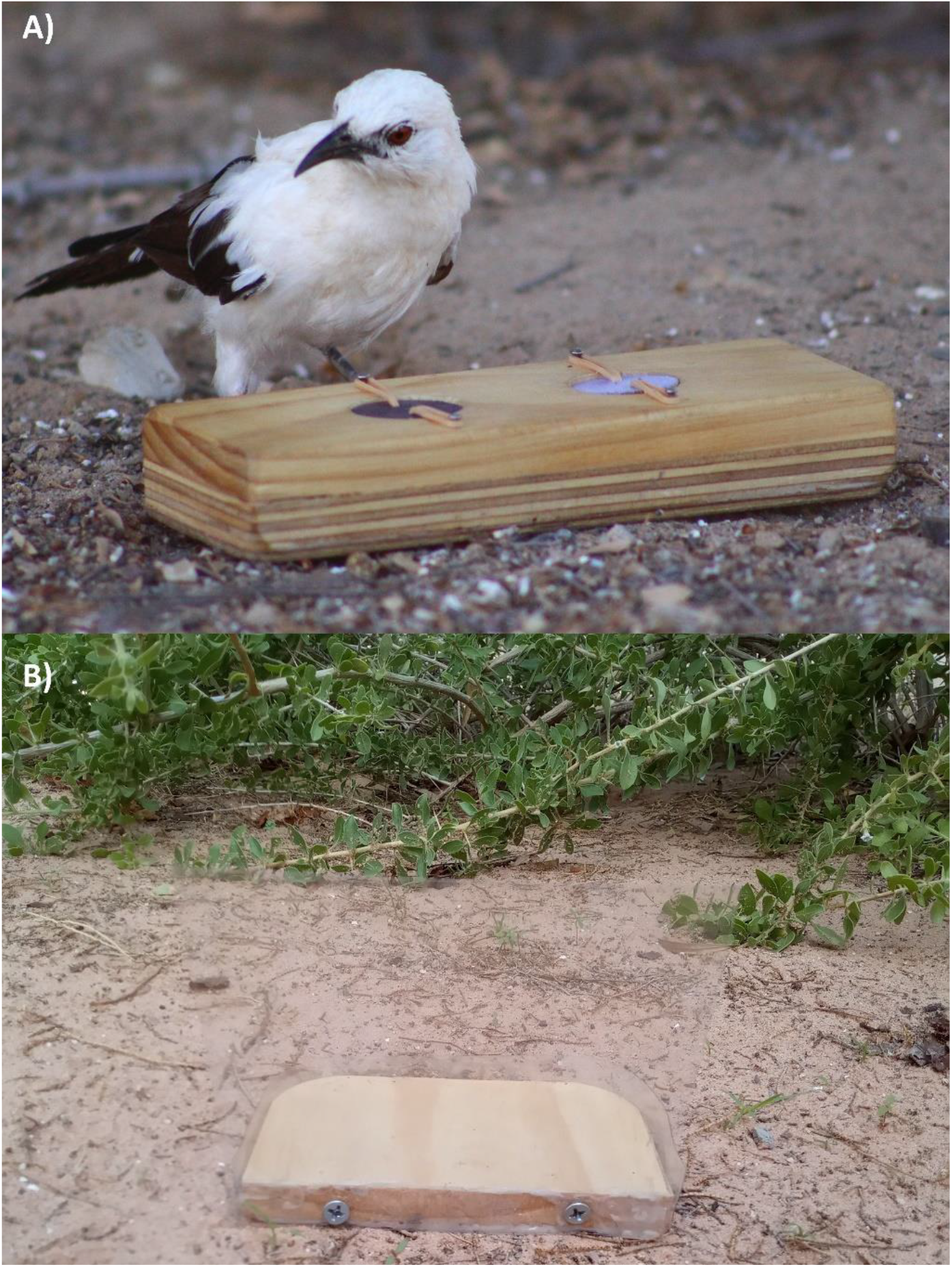
Wild pied babbler interacting with the cognitive task used to quantify associative and reversal learning (A); and example of task used to quantify inhibitory control (B). In A) the individual has to learn the association between a colour cue (dark versus light purple lids in the picture) and a food reward (mealworm inside the well). In B) a mealworm is placed behind the transparent barrier and the individual has to inhibit the prepotent instinct of pecking the barrier when seeing the food reward behind it and instead detour around it. Photo credits: Nicholas Pattinson.

#### 2.2.3 Inhibitory control

We quantified inhibitory control using a detour-reaching task, which consisted of a transparent barrier (clear smooth PVC, 200 µm thick) fixed onto a wooden base (Figure 1B), with a mealworm positioned ∼2 cm behind the barrier on the wooden base. The task was presented to the individual straight on so that the mealworm was visible behind the barrier, but not accessible from the direction that the individual was approaching the task. In this task, the individual had to inhibit the prepotent response of pecking the barrier when seeing the food reward and instead detour around the barrier to retrieve it. A trial was marked as correct if the individual retrieved the mealworm from behind the barrier without pecking it. The passing criterion was six correct trials in a row and the performance measure was the number of trials to criterion.

#### 2.2.4 Task variants

This study was conceived as part of a long-term project that involved repeatedly quantifying individual cognitive performance (see Supplementary Material section 6). To control for the potentially confounding effects of memory on cognitive performance, causally identical but visually distinct variants of each task were used over the course of the project [54]. Different colour combinations (dark vs light green and purple in 2018; dark vs light green, purple, blue, orange, and pink in 2019 and 2021) and shapes of the transparent barrier (cylinder and wall in 2018; cylinder, wall, arch, umbrella, and corner in 2019 and 2021; Figure S1) were randomly assigned to each individual tested. The variant used did not significantly affect the number of trials taken to pass the associative and reversal learning tasks nor the inhibitory control task, respectively (see Supplementary Material section 2).

### 2.3 Proxies of motivation

Performance in cognitive tasks may be influenced by the motivation of the individual to interact with the task, especially in the wild, where researchers have limited control over environmental and individual condition [11]. When completing a cognitive task based on a food reward, individual performance might vary depending on hunger level and amount of food available in the environment. For this reason, we measured several proxies of motivation: foraging efficiency, body mass, latency to approach the task, and inter-trial interval.

Weekly 20-min behavioural focal observations were carried out for all the individuals tested. Focal observations were conducted by continuously recording the behaviours of the individual (to the nearest second) using a customised programme created in the free software Cybertracker. Foraging efficiency [grams of biomass consumed per foraging minute; following 55] was calculated from focal observations comprising at least five minutes of foraging. Food items that were provisioned to young were excluded from the calculation to better approximate individual hunger level. As a previous study found babblers forage more efficiently in the early morning [56], we paired the timing of the focal observation and cognitive testing by performing both either in the early morning (before 9 am) or later in the day (after 9 am). As an additional proxy of hunger level, we measured the body mass of each individual (accuracy 0.1 g) within the four hours prior to each cognitive test by enticing the individual to jump on a top-pan scale to retrieve a mealworm [57]. Finally, we measured the latency to approach the task as the time elapsed between the focal individual being within 5 m of the task and first making contact with the task [5] and the average inter-trial interval (see Supplementary Material section 4).

### 2.4 Measures of reproductive success

Since 2003, each year during the breeding season (September-March) researchers perform weekly visits to the babbler groups during which the number and identity of individuals (adults, fledglings, juveniles) and any breeding activity are noted [44]. Nests are located by observing nest building, and accurate hatch and fledge dates are recorded by checking the nests every two-three days once they have been located. If fledglings are missing after two consecutive visits to the group, they are considered dead. We assumed only dominant individuals bred [46], therefore the offspring produced in each breeding attempt were attributed to the dominant male and female in the group at the time the breeding attempt was recorded. The extensive life history database allowed us to determine the number of fledglings produced per year, the number of fledglings surviving to independence [i.e. 90 days post-hatching, when offspring receive < 1 feed/hour; 57] per year, and the number of fledglings recruited into the adult population [i.e. surviving to one year post-hatching; 58] per year for each dominant individual.

### 2.5 Statistical analyses

All analyses were performed with R statistical software version 4.2.0 [59]. To investigate the causes and fitness consequences of variation in cognitive performance, we fitted different sets of Generalized Linear Mixed Models (GLMMs) using the *lmerTest* package [60] and tested the relative importance of different candidate explanatory terms by ranking them by Akaike information criterion score corrected for small sample sizes (AICc). Models within 2 ΔAICc of the best model and with predictors whose 95% confidence intervals did not intersect zero were included in the top model set, and were considered to explain variation in the dependent variable better than other candidate models [61]. Continuous predictors were scaled by centring on the mean and dividing by one standard deviation. Normality of residuals, presence of outliers and dispersion were checked using the *DHaRMa* package [62].

#### 2.5.1 Relationships between individual cognitive performances across tasks

First, we tested whether cognitive performance was correlated across tasks by performing Spearman’s rank correlations on the scores (i.e. number of trials to pass) of each pair of tasks. Note that a lower score in this case indicates fewer trials to pass the task, and hence, better cognitive performance. To determine whether individual performance in different tasks could be explained by a single factor (GCP), we then performed an unrotated principal component analysis (PCA) on the scores of the associative learning, reversal learning and inhibitory control tasks, using the *FactoMineR* package [63]. Following Shaw et al. [41], to test whether the mean and standard deviation of the loadings onto the first principal component (PC1) deviated from what is expected by chance, we performed 10000 PCA simulations using the function *randomizeMatrix* in the *picante* package [64]. For each simulation, the cognitive scores within each task were randomised among individuals and a PCA was performed. We then compared the real mean and standard deviation of the loadings onto PC1 to the 95% confidence intervals (CI) of the simulated means and standard deviations of the loadings onto PC1.

#### 2.5.2 Factors explaining interindividual variation in cognitive performance

To determine whether individual and group attributes or proxies of motivation explained inter-individual variation in cognitive performance, we fitted LMMs containing group identity as a random term and GCP as dependent variable, where GCP was the individual coordinate along PC1 but with the opposite sign so that higher values corresponded to higher general cognitive performance. We used GCP as a measure of individual cognitive performance because performance on all tasks loaded strongly and positively onto PC1 (section 2.5.1). The individual and group attributes considered as candidate explanatory terms were age, sex, rank, and group size. The proxies of motivation tested were: average latency to approach (s), inter-trial interval (min), body mass (g) and foraging efficiency (g/min), all of which were averaged across the three tasks used to compute GCP. We also included testing order within a group to test for any potential effect of social learning (*sensu* Ashton et al. 2018a). If social learning was occurring, we predicted that individuals tested later within a group would perform better than those tested earlier. Additionally, to test for a potential effect of different activity levels throughout the day, we included the explanatory term “time of day”, which was calculated as follows: each task was assigned a 1 if the test started before 9 am or a 0 if the test started later in the day, then this value was summed for the three tasks, obtaining a value between 0 (all tests started after 9 am) and 3 (all tests started before 9 am). Finally, we included study year (2018; 2019 or 2021) as a predictor to check for any differences in overall conditions across years that might have affected cognitive performance. We also tested all additive models and pairwise interactions among sex, age, rank, group size, body mass and study year. Individuals of unknown sex (N = 6) were excluded from this analysis.

#### 2.5.3 The relationship between cognitive performance and reproductive success

When analysing reproductive success, we considered only dominant individuals because subordinates do not have the opportunity to breed [46]. We included two individuals that were subordinates in the early years of testing but were retested once they gained dominance, for a total of N = 19 dominant individuals. First, we checked whether the individual attributes that determine variation in cognitive performance, i.e. age and sex (based on the results of section 2.5.2), were also associated with variation in the average number of fledglings produced per year since year two of age, which is the earliest age at which individuals in our dataset bred (see Supplementary Material section 8). Hence, we determined if individual cognitive performance was directly related to reproductive success. We considered three measures of reproductive success: number of fledglings produced per year, number of fledglings that survived to independence per year and number of fledglings that survived to recruitment per year. When there were multiple breeding attempts within a year we used cumulative numbers, and we assigned a 0 for years in which dominant individuals did not successfully breed. The average number of years with breeding data per dominant individual tested was 4.7 (range 1-11 years), where a year encompasses the austral breeding season (from September of one year to August of the next year). For each of the three measures of reproductive success we fitted a set of GLMMs with a Poisson error distribution and year and individual ID as random terms. Group ID was not included in these models as a random term because it resulted in overfitting (singular fit). The candidate explanatory terms tested were GCP, age, sex, group size, and drought (1= drought vs 0 = no drought occurring during the breeding season). Group size and drought [defined as rainfall ≤ 137 mm; see 65] were included among the explanatory terms because a recent study found they predicted the number of offspring surviving to independence in babblers [65]. We also tested the interaction between GCP and sex to determine if the relationship between cognition and reproduction differed in males and females. To identify the minimum determinable effect of two-way interactions given our sample sizes [66, 67], we conducted a power analysis with the *pwr* package [68].

## 3. RESULTS

### 3.1 Relationships between individual cognitive performance across tasks

The 38 tested babblers completed the associative learning, reversal learning and inhibitory control tasks in a mean of 39.26 trials (range 6-120), 63.18 trials (range 6-120) and 34.58 trials (range 6-105) respectively; the range indicating great variation in cognitive performance. We found positive correlations in cognitive performance for all pairwise comparisons across tasks, but only the correlation between associative and reversal learning performance was significant (associative and reversal learning: Spearman’s rho = 0.65, p < 0.001; reversal learning and inhibitory control: Spearman’s rho = 0.29, p = 0.08; associative learning and inhibitory control: Spearman’s rho = 0.14, p = 0.42). The consistent positive direction of pairwise correlations between tasks aligns with the output of the PCA, which showed that all cognitive scores loaded positively onto PC1 extracted with an eigenvalue over one (Table 1). PC1 explained 59.5 % of the total variation in cognitive performance across tasks (Table 1).

**Table 1.** Output of the principal component analysis on the scores (i.e. number of trials to pass) obtained by 38 pied babblers on three cognitive tasks quantifying associative learning, reversal learning and inhibitory control.

| Cognitive task | PC1 |
| --- | --- |
| Associative learning | 0.82 |
| Reversal learning | 0.88 |
| Inhibitory control | 0.58 |
| Eigenvalue | 1.79 |
| % Variance explained | 59.54 |

The PCA results were highly unlikely to occur by chance because the real mean loading onto PC1 was higher than the 95% CI of the randomly simulated mean loadings (95% CI of simulated means for PC1 = 0.01-0.67, real mean = 0.76; Figure S3), and while the real SD was within the 95% CI of the simulated SD, it was at the lower end of the distribution (95% CI of simulated SD for PC1 = 0.08-0.83, real SD = 0.16; Figure S3). In other words, of the 10000 random simulations, only 0.03% had a larger mean loading on PC1 and only 8.07% had a smaller SD. Additionally, when we examined the cognitive scores of 18 individuals that were tested twice on the cognitive test battery during the study years (2018-2021), individual scores from the second replicate of the cognitive test battery also loaded positively onto PC1, which explained 46.3% of the total variance in cognitive performance, providing further evidence for general cognitive performance (GCP); importantly, GCP was significantly repeatable (R = 0.50; SE = 0.18; 95% CI = 0.09; 0.78; p = 0.015) (see Supplementary Material section 6).

### 3.2 Factors explaining interindividual variation in cognitive performance

The factors that best explained variation in GCP were age and sex (Table 2). GCP declined with age in females but not males (females: coefficient ± SE = -0.77 ± 0.20, 95% CI = -1.18; -0.37, males: coefficient ± SE = 0.13 ± 0.24, 95% CI = -0.36; 0.62; N = 32, of which 16 females and 16 males; see Figure 2). Group size was not a significant predictor of general cognitive performance (see Supplementary Material section 9 for a discussion of this result). Importantly, study year and the proxies of motivation examined (i.e. latency to approach the task, inter-trial interval, body mass, foraging efficiency, time of day) did not significantly explain variation in GCP (Supplementary Material Table S3).

**Table 2.**
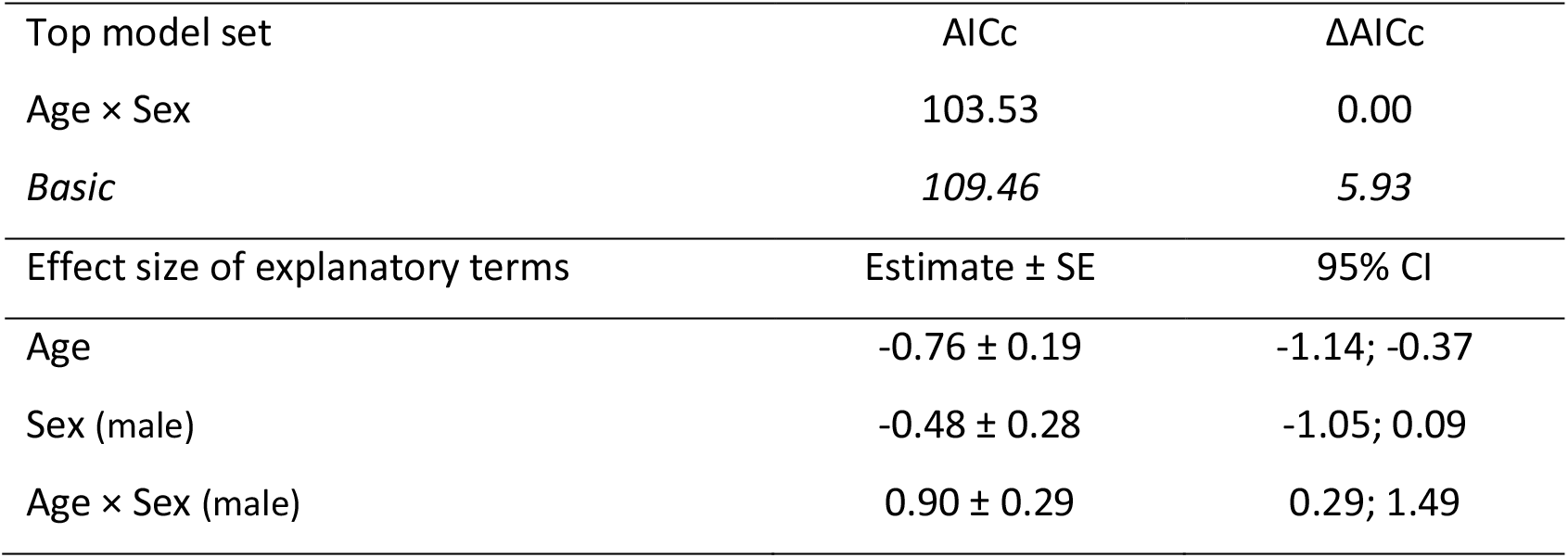
Top model set of candidate terms affecting general cognitive performance in pied babblers. All models included group ID as a random term. Corrected Akaike information criterion (AICc) and ΔAICc are provided for models within 2 ΔAICc of the top model and with predictors whose 95% confidence intervals (CI) do not intersect zero. Coefficient estimates ± standard errors (SE) and 95% CI are given below the top model set. N = 32 individuals from 11 groups. See Supplementary Material, Table S3 for full model selection outputs.

**Figure 2.**
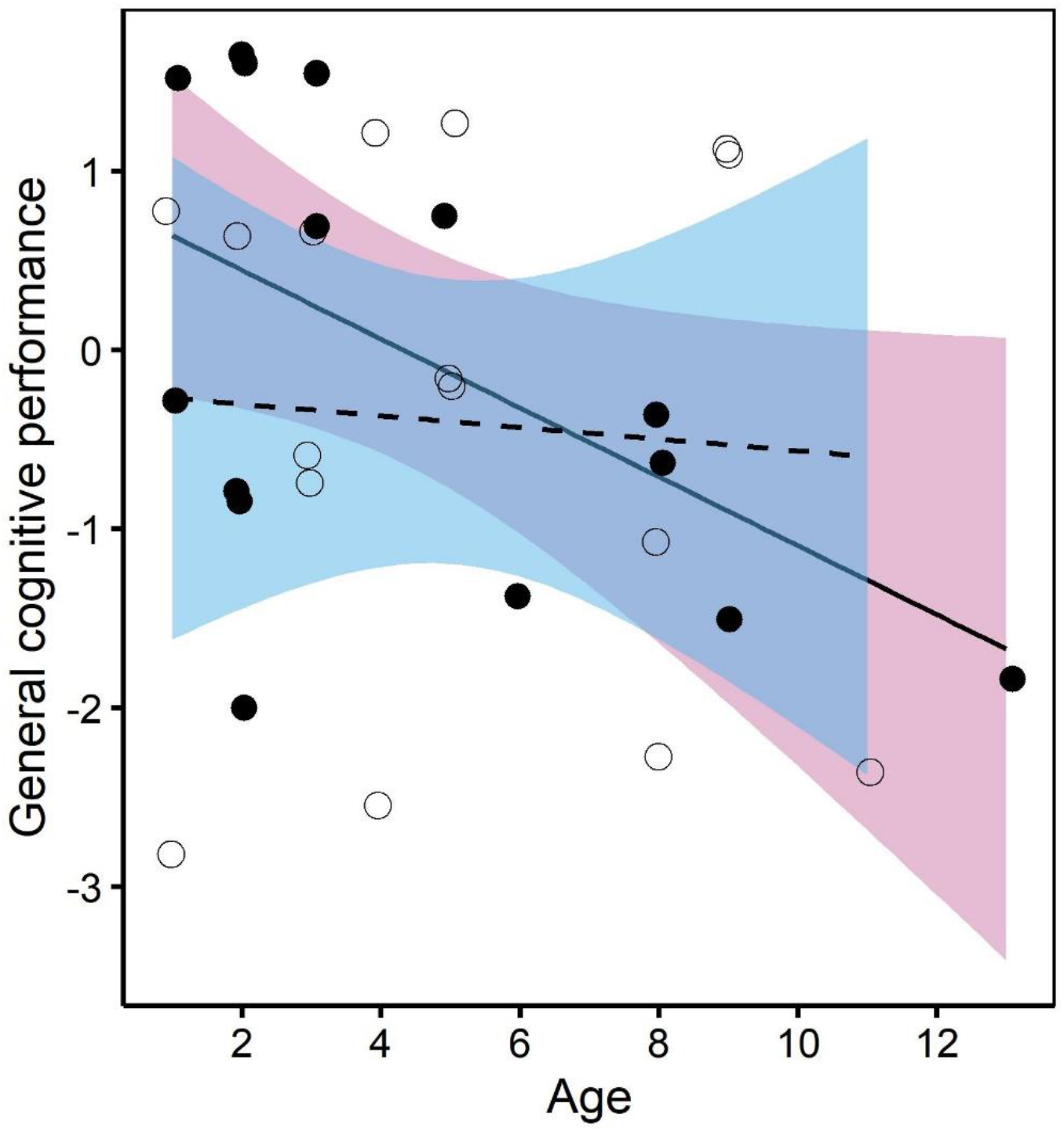
Variation in pied babblers’ general cognitive performance by age and sex (females: pink colour, solid line, filled dots; males: blue colour, dashed line, empty dots). General cognitive performance declined with age in females but not in males (N = 16 females and 16 males). Points are raw data; fitted lines and 95% confidence interval bands are generated from the output of the model presented in Table 2.

### 3.3 The relationship between general cognitive performance and reproductive success

The average number of fledglings produced per year since age two tended to increase with age in females but not in males (Supplementary Material section 8). Hence, in females, the relationship between reproductive success and age followed an opposite trend compared to the relationship between GCP and age: older females tended to produce more fledglings per year on average but showed lower general cognitive performance. In line with this result, we found that individual general cognitive performance was negatively related with the number of fledglings produced per year (Table 3A, Figure 3). We did not find evidence that this relationship differed in males and females (non-significant interaction GCP × sex), but we only had power to detect very large effects of two-way interactions (Cohen’s f^2^ = 0.51).

**Table 3.** Model set of candidate terms affecting three measures of reproductive success in pied babblers. The models included year and individual ID as random terms. Corrected Akaike information criterion (AICc) and ΔAICc are provided for candidate explanatory terms. Coefficient estimates ± standard errors (SE) and 95% confidence intervals (CI) are given below the model sets for models within 2 ΔAICc of the top model and with predictors whose 95% confidence intervals (CI) do not intersect zero. The measures of reproductive success examined were (A) number of fledglings produced per year, N = 90 observations for 19 dominant individuals over 14 years; (B) number of fledglings reaching independence per year, N = 81 observations for 18 dominant individuals over 14 years; (C) number of fledglings recruited per year, N = 79 observations for 14 dominant individuals over 13 years.

| Model selection | AICc | ΔAICc |
| --- | --- | --- |
| A) Number of fledglings per year |  |  |
| General cognitive performance | 356.08 | 0.00 |
| General cognitive performance × Sex* | 357.38 | 1.30 |
| Sex* | 357.46 | 1.38 |
| Age | 358.76 | 2.68 |
| Basic | 359.01 | 2.93 |
| Drought | 359.75 | 3.67 |
| Group size | 361.17 | 5.09 |
| B) Number of independent offspring per year |  |  |
| Drought | 258.70 | 0.00 |
| Basic | 264.59 | 5.89 |
| Sex | 264.80 | 6.10 |
| Age | 266.09 | 7.39 |
| General cognitive performance | 266.26 | 7.56 |
| Group size | 266.56 | 7.86 |
| General cognitive performance × Sex | 269.21 | 10.51 |
| C) Number of offspring recruited per year |  |  |
| Drought | 204.48 | 0.00 |
| Basic | 205.73 | 1.25 |
| Age | 205.77 | 1.29 |
| Sex | 206.05 | 1.57 |
| General cognitive performance | 206.74 | 2.26 |
| Group size | 207.80 | 3.32 |
| General cognitive performance × Sex | 209.27 | 4.79 |
| Effect size of explanatory terms | Estimate ± SE | 95% CI |
| A) Number of fledglings per year |  |  |
| General cognitive performance | -0.18 ± 0.07 | -0.34; -0.03 |
| B) Number of independent offspring per year |  |  |
| Drought | -0.99 ± 0.34 | -1.82; -0.36 |

\* Not included in the top model set because 95% CI intersect zero

**Figure 3.**
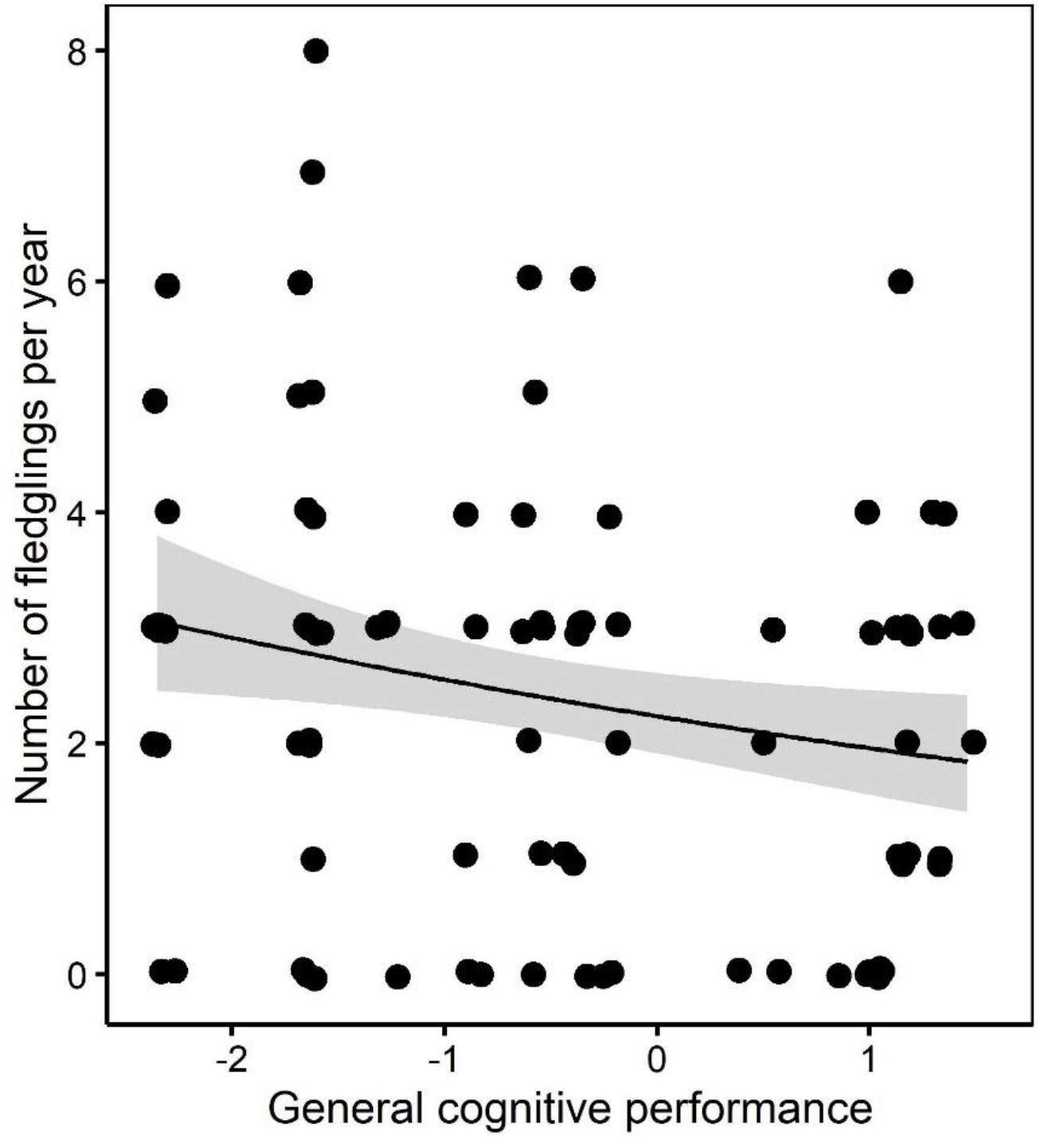
The relationship between the number of fledglings produced per year and general cognitive performance in dominant pied babblers (N = 90 observations for 19 dominant individuals over 14 years). Individuals showing higher general cognitive performance produced fewer fledglings per year. Points are raw data; the fitted line and 95% confidence interval band are generated from the output of the model presented in Table 3.

The main predictor of the number of fledglings surviving to independence was the occurrence of droughts, with more fledglings reaching nutritional independence in non-drought years (Table 3B). None of the explanatory terms tested were a significant predictor of the number of fledglings surviving to recruitment (Table 3C). However, we had to exclude the two dominant females who showed the highest GCP from the latter analysis due to missing data on the number of fledglings surviving to recruitment, therefore the lack of an effect of GCP on the number of offspring recruited per year should be interpreted with caution.

## 4. DISCUSSION

We quantified individual cognitive performance in a wild bird population with the aim to answer three central questions in cognitive ecology: (a) does performance co-vary across cognitive tasks, (b) what drives these individual differences, and (c) is individual cognitive performance related to reproductive success. We found that most of the variation in individual cognitive performance across tasks could be explained by a single factor (GCP or general cognitive performance). Individual differences in GCP depended on age and sex. Older females (but not males) showed lower GCP and tended to produce more fledglings per year on average. Accordingly, we found that GCP was negatively related to the number of fledglings produced per year. These findings support the existence of general cognitive processes in wild babblers and suggest that individuals might trade off the resources invested in reproduction against cognitive performance.

### 4.1 Is there evidence for a general cognitive factor in wild babblers?

Individuals that learnt an association faster, were also faster at reversing the learnt association and showed better inhibitory control, as indicated by positive (albeit not always significant) correlations in cognitive performance across tasks. Indeed, approximately 60% of the variance in individual cognitive performance across tasks could be explained by a single factor: GCP. Additionally, GCP was significantly repeatable (R = 0.50), indicating that our measure of general cognitive performance captured consistent inter-individual differences in cognition. While we cannot completely exclude that motivation to interact with the cognitive tasks affected cognitive performance, we are confident that its effect on our measure of GCP was minimal because none of the measured proxies of motivation, such as average latency to approach the task or inter-trial interval, significantly explained variation in GCP. It is also worth noting that all the tested birds interacted with the tasks and always ate the food, further indicating that the birds were motivated to interact with the tasks. Therefore, our findings are consistent with the existence of a general cognitive factor underpinning performance across different cognitive domains in babblers. However, our test battery included only three cognitive tasks, which is the minimum number required to test for a general cognitive factor [35]. Therefore, future studies should consider expanding the test battery by including, for example, spatial memory tasks redesigned so that there is scope to quantify spatial memory [e.g. adding a presentation at 72h or changing the scale of the spatial task; 41], and tasks assessing social cognition, the ability to make inferences, and reaction time [37, 41].

Alternative explanations for the single factor GCP underpinning individual cognitive performance across tasks are also possible. First, the different tasks used may tap into the same cognitive process. For example, it has been suggested that associative learning may underlie variation in performance in animal test batteries [40]. Second, positive correlations between tasks could be the consequence of underlying variation in individual phenotypic or genetic quality; for example, a single genetically-determined component of the nervous system may determine differences in neuronal function that affect all cognitive domains simultaneously [69]. Therefore, whether statistical evidence for GCP indicates a truly general cognitive ability underlying performance across different cognitive domains remains to be determined.

### 4.2 Age-related cognitive decline and individual reproductive success

In babblers, individual variation in general cognitive performance was predicted by an interaction between age and sex, with cognitive performance declining with age in females but not in males. Faster cognitive ageing in females has been previously reported in humans [70], nematodes (*Caenorhabditis remanei*) [71], mice (*Mus musculus*) [72], and captive marmosets (*Callithrix jacchus*) [25]. However, the only study testing for cognitive senescence in the wild found no decline in spatial memory performance in mountain chickadees (*Poecile gambeli*) from one to six years of age [26]. Hence, to our knowledge, our finding represents the first evidence of sex differences in age-related cognitive decline in a wild animal.

Senescence has been explained by two main evolutionary theories [reviewed in 28]. The “selection shadow” theory states that selection strength decreases with age after sexual maturity [73]. Our data do not support this theory because babblers were still breeding up to 13 years of age, leaving ample opportunity for selection to act on cognitive traits among older individuals. A second theory is the life history theory of ageing, which encompasses two convergent theories: the first states that due to the limited resources available to organisms, these must be traded-off between reproduction and somatic maintenance (“disposable soma theory”) [74]; the second states that alleles with beneficial effects early in life but detrimental effects later in life can be favoured by selection (“antagonistic pleiotropy”) [75]. Based on these theories we would expect that in babblers (1) cognitive performance in older females is traded-off against increased reproductive output, and/or that (2) females have been selected for higher cognitive performance early in life even at the expenses of reduced cognitive performance later in life.

Previous studies on babbler life history [51, 76] provide some support for both explanation (1) and (2), which are not mutually exclusive. First, female babblers (but not males) engage in costly breeding competition [76, 77]. Subordinate females compete both indirectly, by courting and nest-building with unrelated dominant males, and directly, by destroying the eggs of the dominant female [76]. This competition forces dominant (and older) females to engage in frequent aggressive displays towards subordinate (and younger) females and repeatedly abandon breeding attempts and re-lay clutches [76], which entails an additional energetic cost [78]. Hence, in older (dominant) females the cost of maintaining a high reproductive output, even in the presence of competitors, might be traded-off against the maintenance of the energetically costly nervous system [13]. For example, previous experiments in the fruit fly and the cabbage white butterfly (*Pieris rapae*) have revealed a trade-off between learning performance and competitive ability [79] or female fecundity [80], respectively. In line with this explanation, when analysing long-term reproductive success in dominant babblers, we found that higher cognitive performance was associated with a lower number of fledglings produced per year. However, a larger sample size will be necessary to test whether this negative relationship between cognition and reproduction differs between males and females.

The second explanation (i.e. selection for higher female cognitive performance earlier in life) is partly supported by sex differences in babbler dispersal strategies. Females are more likely than males to gain a breeding position by overthrowing a dominant female in a non-natal group [51, 81]. Accordingly, juvenile females are more aggressive than males, and higher female aggressiveness is associated with younger age at dispersal [82]. On the contrary, males are more sedentary [83] and disperse only when search costs are low [84]. It is possible that these sex differences lead to selection on females for higher cognitive performance early in life, even at the expenses of faster cognitive senescence. Indeed, cognitive performance in young females might be crucial to gain access to breeding positions by enabling them to navigate across territories, identify the sex and rank of conspecifics in non-natal groups, and decide when to engage in aggressive displays towards a dominant female [85]. Additionally, in babblers the number of immigrant competitors decreases with pair bond tenure, while reproductive success increases [48]. This suggests that on average the risk of losing a breeding position and thus, potentially, the need to maintain high cognitive performance might decrease with female age. Overall, our findings paired with evidence from previous research in babblers suggest that females may be under selection for higher cognitive performance earlier in life despite faster cognitive senescence and/or cognitive senescence may be accelerated by investment in reproduction and breeding competition. However, longitudinal studies are ultimately needed to describe cognitive ageing trajectories and test these hypotheses.

Since we used a cross-sectional design instead of a longitudinal design, we cannot determine whether cognitive performance declined throughout life in females, or whether only females with lower cognitive performance survived until old ages. Therefore, a third potential explanation for the observed sex differences in age-related cognitive decline is that cognitive performance is negatively linked to survival, at least in females. For example in pheasants (*Phasianus colchicus*), survival in the wild was negatively related to reversal learning performance [86]. As most of the birds tested in the present study are still alive to date, we could not perform a survival analysis to examine potential effects of individual cognitive performance on survival, but this will be a necessary next step to confirm whether the findings from the present study are due to cognitive senescence or reduced survival of smarter females.

### 4.3 A negative relationship between cognitive performance and reproductive success

We found that individuals with better general cognitive performance produced fewer fledglings per year, which is consistent with a trade-off between cognition and reproduction. However, GCP did not predict the number of fledglings surviving to nutritional independence, which depended instead on the occurrence of droughts during the breeding season, in line with a previous study [65]. It is possible that parental traits influence offspring survival in the nestling stage but not in the post-fledgling stage, where survival may be more strongly influenced by environmental conditions [87]. Therefore, the extent to which cognitive performance may be under negative selection remains to be determined. Cognitive performance might also be simultaneously associated with life-history traits linked to fitness in different directions [88, 89]. For example, while cognitive performance is negatively related to the number of fledglings produced per year by dominant individuals, it might be positively related to the age at which dominance is acquired in the first place. Future comparisons of the age at acquisition of dominance among individuals whose cognition was tested as subordinates will allow us to address this hypothesis.

### 4.4 Conclusion

We found that individual cognitive performance covaried across tasks, which is consistent with a general cognitive factor, though alternative explanations cannot be excluded. We considered the effect of individual and social attributes and several proxies of motivation on cognitive performance. We found that general cognitive performance depended on sex and age, declining with age in females but not males. Older females also tended to fledge more nestlings per year. By analysing over 10 years of breeding data, we show that individuals with lower general cognitive performance produced more fledglings per year. Our findings suggest that cognitive performance is traded-off against reproduction, demonstrating that in order to understand how selection acts on cognition we need to consider not only its benefits but also its costs.

## Acknowledgements

We thank the managers at the Kuruman River Reserve (KRR) and the families at the surrounding farms (Van Zylsrus, South Africa) for making this work possible. We thank Amanda Bourne, Lina Peña-Ramirez, Grace Blackburn, Amy Hunter, and Samantha Wagstaff for assistance during data collection. We are grateful to the Hot Birds Research Project for access to the KRR rainfall data and to Amanda Bourne for calculation of drought/non-drought years. We thank the FitzPatrick Institute of African Ornithology for ongoing logistical support. We also thank Prof Leigh Simmons for the advice given to Camilla Soravia as part of her supervisory team.

## Ethics statement

This research was approved by the Animal Ethics Committee, University of Western Australia (RA/3/100/1619 and 2021/ET000414).

