## Supplementary Material for "General cognitive performance declines with female age and is negatively related to fledging success in a wild bird"

### 1. Training

Before cognitive testing, we trained the birds to peck the lids in a cognitive task to find a food reward. The array used for training was the one used in the associative/reversal learning task (small wooden block measuring 180 x 70 x 30 mm with two circular wells of 30 mm diameter and 20 mm depth covered by wooden lids). During training, both wells were baited and both wooden lids had a black cross drawn over the wooden background. This prevented the birds from forming any association between a specific colour and the food reward prior to cognitive testing. The training followed a shaping procedure in four steps [1-3]: the task was presented (1) without lids; (2) with the lids on the array but not covering the wells; (3) with the lids partially covering the wells; and (4) with the lids fully covering the wells. If the bird retrieved the mealworm in one step, training moved to the following step, if the bird did not retrieve the mealworm for three presentations in a row, it went back to the previous step. Training was completed when the bird retrieved the mealworm from fully covered wells six times in a row, which represents a significant deviation from a random binomial probability. Cognitive testing was carried out at least 24h after training was completed.

### 2. Task variants

For the purposes of another study, we planned to quantify cognitive traits multiple times. To avoid confounding effects of memory across replicates, different task variants had to be used, i.e. versions of a task that differ in their visual appearance but maintain the exact same functioning [2, 4]. The variants used for the associative and reversal learning tasks comprised a dark and light shade of one of the following colours: green, blue, purple, orange, pink. There was no difference in the number of trials required to reach learning criterion or the upper cut-off of 120 trials depending on variant (associative learning: Kruskal-Wallis test $\chi^2(9) = 9.50$, $p = 0.392$; reversal learning: Kruskal-Wallis test $\chi^2(9) = 14.60$, $p = 0.103$). Similarly, the variants used for the inhibitory control task consisted of different shaped barriers: wall, corner, arch, umbrella, cylinder (see Figure S1). There was no difference in the number of trials required to pass the inhibitory control task depending on variant (Kruskal-Wallis test $\chi^2(4) = 1.34$, $p = 0.855$).

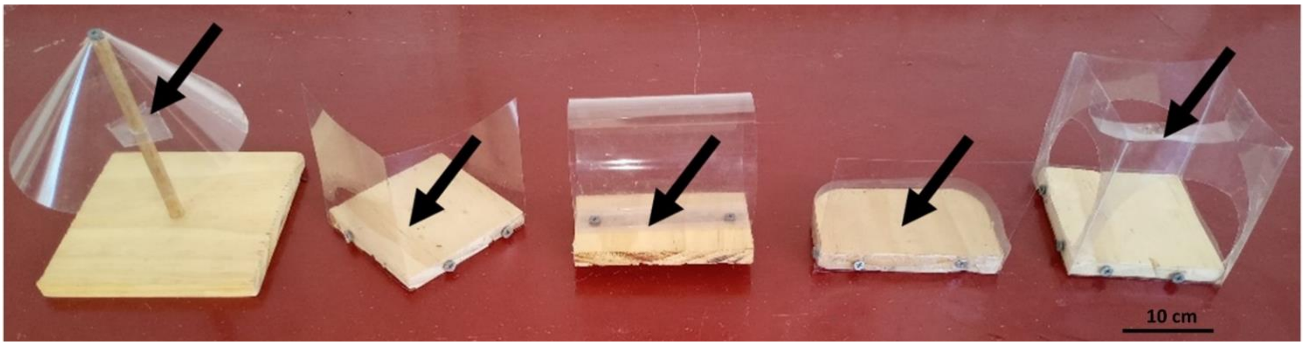

**Figure S1.** Variants of the transparent barrier used to quantify inhibitory control; from left to right: umbrella, corner, cylinder, wall, arch. The black arrows indicate where the mealworm (*Tenebrio molitor* larva) was positioned. In all variants, the mealworm appeared behind the transparent PVC barrier from the perspective of an approaching pied babbler on the ground. To complete a trial correctly an individual had to inhibit the prepotent instinct of pecking the transparent barrier when approaching the task and instead detour to retrieve the food reward. The inhibitory control tasks required different routes of detour; in the corner, wall, and cylinder an individual had to detour towards the open ends of the barrier; whereas in the umbrella an individual had to detour underneath the barrier, and in the arch an individual could detour either underneath or above the barrier.

#### 3. Spatial memory task

The spatial memory task consisted of a wooden foraging grid with eight equidistant wells (in three rows of two, four, two wells respectively; Figure S2) of the same shape and depth as the ones used in the associative and reversal learning task. All wells were covered by lids of the same shade of grey. The food reward consisted of two mealworms. The rewarded location was randomly assigned to one of these wells for each test bird. The test took place over three days: on the first day, the task was presented to the bird twice (with a five-minute interval), allowing it to search the grid until the rewarded well was found both times; the task was then presented once 24h and once 48h afterwards. The cumulative number of wells searched before locating the rewarded well on the 24h and 48h presentations represented the spatial memory score. Five minutes after the 48h presentation, a control trial for olfactory cues was carried out. In this trial the foraging grid was presented rotated 180 degrees and without a food reward: if the test subject remembered the location of the rewarded well it would search the correct location (now opposite to the previously rewarded well), while if it had been using olfactory cues it would search the previously rewarded well.

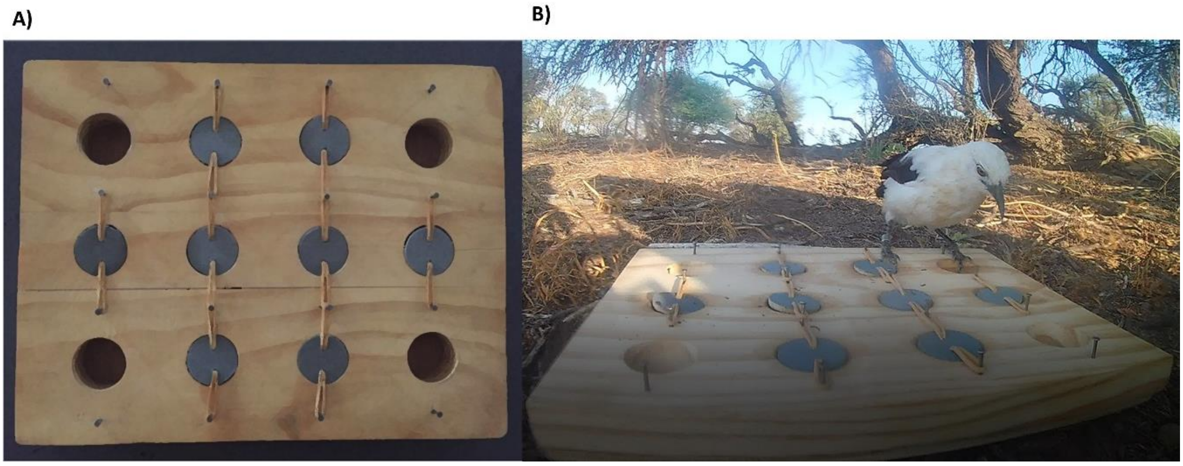


**Figure S2.** Cognitive task used to quantify spatial memory (A); and individual pied babbler interacting with the task in the wild (B). The task consisted of eight equidistant wells covered by grey lids that swivel when pecked (corner wells not used). The food reward (2 mealworms) was randomly assigned to one of the eight wells.

The number of wells that pied babblers were expected to search under a random sampling strategy was calculated based on the equation given by Tillé Y, Newman JAHealy SD [5] for a spatial memory task in which animals are allowed to sample locations until they achieve a fix number of successes and sampling occurs without replacement. In this case the random expectation follows a negative hypergeometric distribution, with the equation $E(Y) = \frac{r(N+1)}{A+1}$, where $r$ = number of successes, $A$ = number of locations that actually contain the food reward, $N$ = total number of locations available to search. In the spatial memory task used here: $r$ = 1, $A$ = 1, $N$ = 8. Therefore, $E(Y)$ = 9/2 = 4.5 wells. Hence if the tested pied babblers searched more than 4.5 wells in the 24h or 48h presentations, they were not deviating from a random sampling strategy.

Pied babblers (N = 33) made between 3-15 (median 9) cumulative mistakes in total across the 24h and 48h presentations. Pied babblers did not search significantly less than the random expectation of 9 when considering the cumulative number of wells searched across the 24h and 48h presentations (Wilcoxon signed rank test: V = 154, p = 0.295). In either presentation, pied babblers did not search significantly less wells than the random expectation of 4.5 (24h presentation: median 5, range 1-8, Wilcoxon signed rank test: V = 272, p = 0.443; 48h presentation: median 4, range 2-8, Wilcoxon signed rank test: V = 219, p = 0.134).

### 4. Proxies of motivation: latency to approach the task and inter-trial interval

During each trial of a cognitive test, the experimenter measured the latency to approach the task as the time elapsed between the focal individual being within 5m of the task and first making contact with the task [2]. Latency to approach the task was then averaged across all trials of a given cognitive test to obtain a proxy of overall motivation. Time was also noted down for each trial, allowing us to estimate the average inter-trial interval for a given cognitive test. If an individual was temporarily out of sight due to incubation or the group moving to a different foraging area, time was discarded from the calculation of the inter-trial interval. Hence, average inter-trial interval is a measure of how long the individual took on average to approach the task for a second time after the previous trial when the task was available to the individual and external factors such as group movements were not influencing bird behaviour.

### 5. Output of 10000 PCA on cognitive scores randomised among individuals

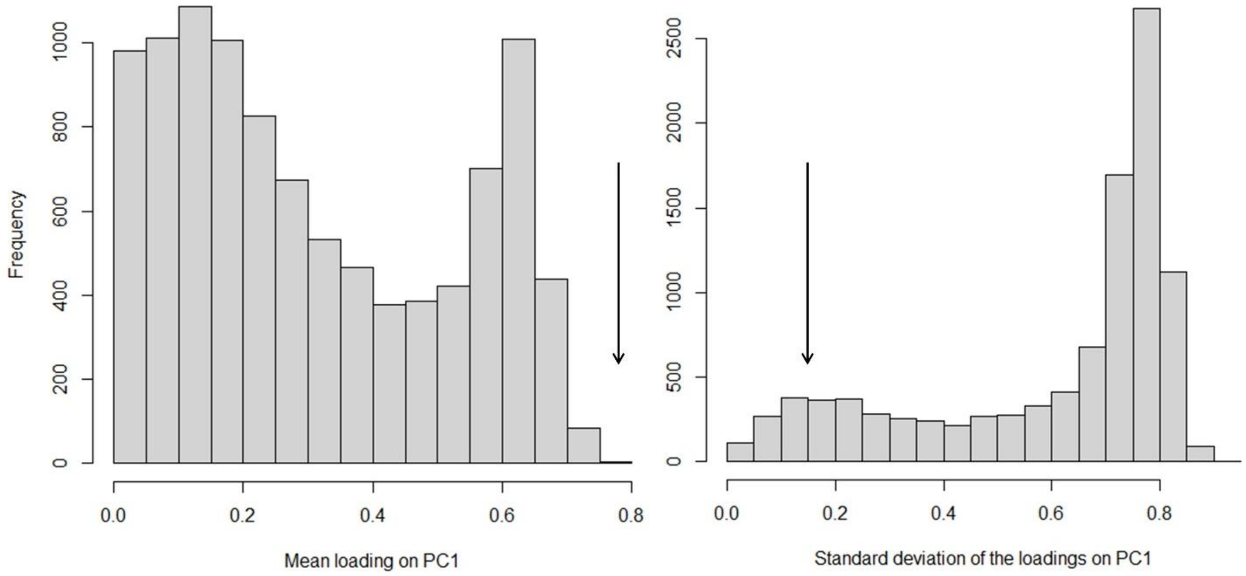


**Figure S3.** Histograms of the means (left panel) and standard deviations (right panel) of the cognitive task loadings onto PC1 generated from 10000 PCA simulations on cognitive scores randomised among individuals. The real mean (0.76) and standard deviation (0.16) of cognitive task loadings onto PC1 are marked in the plot by arrows. Only 0.03% of the simulations had a larger mean than the one obtained from real data and only 8.07% of the simulations had a lower standard deviation, indicating that the pattern of cognitive task loadings onto PC1 found in this study is unlikely to occur by chance.

### 6. Repeatability estimates

In order to test how reliable measures of individual cognitive performance are, we calculated repeatability estimates using the R package *RptR* [6]. First, we estimated repeatability separately for each cognitive task (i.e. associative learning, reversal learning, inhibitory control). To this aim, we fitted Generalised Linear Mixed Models (GLMMs) with Poisson distribution, setting cognitive performance (i.e. number of trials to pass) as the dependent variable, and individual ID as a random term. Some individuals were tested three times on a given task over the study period (2018-2021); in order to increase the statistical power for the repeatability analysis, we included cognitive performance measures from all three replicates for these individuals. Therefore, the final sample sizes were: associative learning (N = 53 tests from 23 individuals of which 7 tested 3 times and 16 tested 2 times); reversal learning (N = 43 tests from 19 individuals of which 5 tested 3 times and 14 tested 2 times); inhibitory control (N = 56 tests from 24 individuals of which 8 tested 3 times and 16 tested 2 times).
Second, we estimated repeatability of general cognitive performance. In this case, we used only the first and second replicate of the cognitive test battery for a given individual. The sample size for this analysis was N = 18 individuals tested twice on all three cognitive tasks. The average time difference between the first two replicates was 330 ± 311 days (SD); the range in time is due to the fact that 8 individuals were retested within the same breeding season and 10 individuals in following seasons. To calculate values of general cognitive performance, we performed two separate principal component analyses (PCAs) on individual cognitive scores in the first and the second replicate of the cognitive test battery using the R package *FactoMineR* [7]. In both replicates, cognitive scores from all three cognitive tasks loaded positively onto PC1, and PC1 had an eigenvalue >1 (see Table S1). Therefore, we extracted individual coordinates along PC1 with the opposite sign so that higher values corresponded to better cognitive performance (i.e. less trials to pass the tasks). Hence, we estimated the repeatability of general cognitive performance by fitting a LMM with individual identity as a random term.

We found that over the course of this study (austral summers 2018-2021) both associative and reversal learning performance were significantly repeatable (Table S2), while inhibitory control performance was not repeatable (Table S2). However, general cognitive performance was significantly repeatable (Table S2).

**Table S1.** Output of the principal component analysis on the scores (i.e. number of trials to pass) obtained by 18 wild adult pied babblers tested twice on three cognitive tasks quantifying associative learning, reversal learning and inhibitory control.

| First Replicate | PC1 | Second Replicate | PC1 |
|---|---|---|---|
| Associative learning | 0.73 | Associative learning | 0.34 |
| Reversal learning | 0.93 | Reversal learning | 0.83 |
| Inhibitory control | 0.64 | Inhibitory control | 0.77 |
| Eigenvalue | 1.80 | Eigenvalue | 1.39 |
| % Variance explained | 60.10 | % Variance explained | 46.26 |

**Table S2.** Repeatability estimates for performance in three cognitive tasks separately and for general cognitive performance (GCP) in wild pied babblers.

| Cognitive measure | R | SE | 95% CI | p |
|---|---|---|---|---|
| Associative learning | 0.30 | 0.14 | 0.01; 0.56 | 0.008 |
| Reversal learning | 0.41 | 0.15 | 0.06; 0.65 | 0.003 |
| Inhibitory control | 0.15 | 0.13 | 0; 0.44 | 0.149 |
| General cognitive performance | 0.50 | 0.18 | 0.09; 0.78 | 0.015 |

### 7. Factors explaining interindividual variation in cognitive performance

**Table S3**. Full model selection output for candidate terms affecting general cognitive performance (GCP) in wild pied babblers. All models included group ID as a random term. Corrected Akaike information criterion (AICc) and ΔAICc are provided for each candidate model. Models within 2 ΔAICc of the top model and with predictors whose 95% confidence intervals (CI) do not intersect zero have been highlighted in bold. The sample size was N = 32 individuals, 11 groups.

| Predictor | AICc | ΔAICc |
|---|---|---|
| **Age × Sex** | **103.53** | **0.00** |
| Age | 107.47 | 3.94 |
| Rank × Sex | 108.02 | 4.49 |
| Age + Sex | 108.22 | 4.69 |
| *Basic* | *109.46* | *5.93* |
| Sex | 110.04 | 6.51 |
| Age + Group size | 110.11 | 6.58 |
| Group size × Year | 110.17 | 6.64 |
| Age + Weight | 110.28 | 6.75 |
| Age × Group size | 110.59 | 7.06 |
| Body mass | 111.03 | 7.50 |
| Rank | 111.31 | 7.78 |
| Testing order | 111.32 | 7.79 |
| Time of day | 111.80 | 8.27 |
| Inter-trial interval | 111.83 | 8.31 |
| Group size | 111.89 | 8.37 |
| Foraging efficiency | 111.94 | 8.41 |
| Latency | 112.00 | 8.48 |
| Body mass + Sex | 112.29 | 8.76 |
| Age + Year | 112.49 | 8.97 |
| Rank + Sex | 112.58 | 9.05 |
| Group size + Sex | 112.77 | 9.24 |
| Body mass × Sex | 112.97 | 9.45 |
| Age × Body mass | 113.24 | 9.71 |
| Group size + Body mass | 113.64 | 10.11 |
| Sex + Year | 113.74 | 10.21 |
| Rank + Group size | 113.85 | 10.32 |
| Age × Year | 113.90 | 10.37 |
| Year | 113.97 | 10.44 |
| Group size × Sex | 114.22 | 10.69 |
| Rank + Year | 115.95 | 12.42 |
| Body mass + Year | 116.23 | 12.70 |
| Group size × Rank | 116.39 | 12.86 |
| Group size × Body mass | 116.42 | 12.89 |
| Sex × Year | 116.44 | 12.91 |
| Group size + Year | 116.76 | 13.23 |
| Rank × Year | 118.22 | 14.69 |
| Body mass × Year | 118.79 | 15.26 |

### 8. Variation in reproductive success by age and sex

We tested whether individual age and sex explained variation in the average number of fledglings produced per year of life, which was calculated as follows: for each dominant individual, we averaged the number of fledglings produced in each year starting from year 2 of age (i.e. earliest age at which individuals in our dataset bred) up to the testing year, assigning a 0 to years in which the individual did not raise any fledglings. The small sample size (N = 19 dominant individuals (≥ 2 years old) from 10 groups, of which 12 males and 7 females) meant that there was only one dominant individual tested for some groups, which resulted in a singular correlation matrix when fitting a LMM with group identity as a random term. Therefore, we fitted a linear model with average number of fledglings produced per year as the dependent variable and the two-way interaction between age and sex as predictor. We found that older females, but not males, produced on average more fledglings per year of life since age 2 (females: coefficient ± SE = 0.96 ± 0.36, 95% CI = 0.19; 1.72, males: coefficient ± SE = -0.05 ± 0.32, 95% CI = -0.72; 0.62, Figure S4). However, the AICc of the model was not 2ΔAICc lower than the intercept-only model (age × sex AICc = 61.47, basic AICc = 62.26); therefore, this result should be treated with caution.

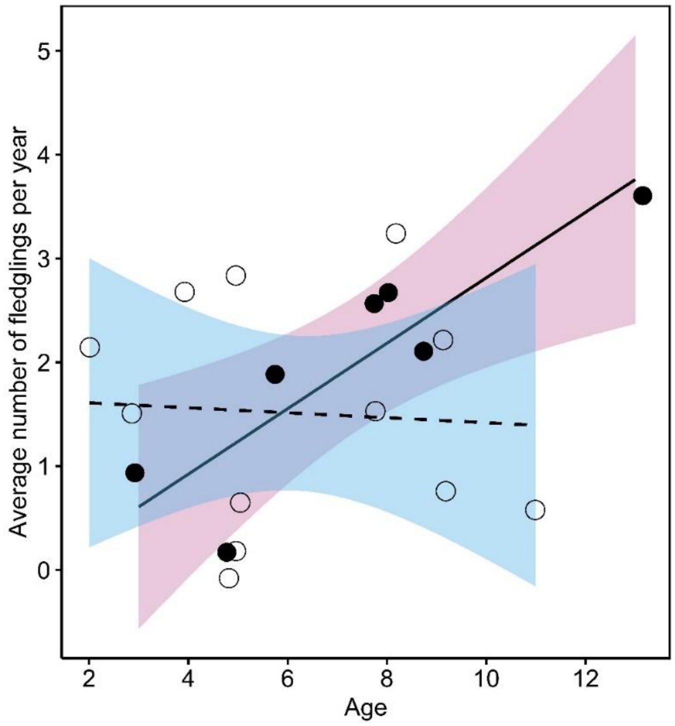


**Figure S4.** Variation in pied babblers' average number of fledglings produced per year by age and sex (females: pink colour, solid line, filled dots; males: blue colour, dashed line, empty dots). The average number of fledglings produced per year tended to increase with age in females but was independent of age in males. Sample sizes: N = 19 dominant individuals (≥ 2 years old) from 10 groups, of which 12 males and 7 females. Points are raw data; fitted lines and 95% confidence interval bands are generated from the output of the model presented in the above text.

### 9. Individual general cognitive performance is independent of group size in pied babblers

We found no evidence that living in larger groups is linked to cognitive performance in pied babblers. This is in contrast with recent findings on Western Australian magpies (*Gymnorhina tibicen dorsalis*), where individuals living in larger groups showed higher GCP [2]. The difference between the two species could depend on the degree to which their societies involve individualised relationships [8]. Babblers live in groups with high within-group relatedness and high reproductive skew [9, 10]. In contrast, in groups of Western Australian magpies, multiple individuals breed within the group, offspring care by helpers is facultative [11] and within-group relatedness is low [12]. These conditions give more scope for conflicts of interest and hence for a society where individuals may benefit from tracking the outcome of past interactions with other group members and negotiate access to resources and breeding opportunities [8, 13]. Therefore, rather than social living *per se*, we must consider the extent to which social systems present individuals with information-processing challenges, such as the need to navigate multiple differentiated social relationships [14]. To test this, future cognition studies should quantify the amount and diversity of social interactions among group members through social network analysis.
